# Differential control of mycobacteria among COVID-19 patients is associated with CD28+ CD8+ T cells

**DOI:** 10.1101/2024.12.09.626962

**Authors:** Alba Llibre, Henna Siddiqui, Jamie Pillaye, Julie G Burel, Charlotte Jones, Harriet Hill, Sian E Faustini, Ella Windle, Hanfa Karim, Emma Sherry, Christopher A Green, Martin Dedicoat, Zania Stamataki, Adam F Cunningham, Matthew K O’Shea

**Author notes:** **Corresponding authors** Matthew K O’Shea Adam Cunningham Alba Llibre.

## Abstract

Diseases caused by SARS-CoV-2 and *Mycobacterium tuberculosis* (*M.tb*) represent two public health emergencies. In severe disease, both pathogens may share a biological niche in the lower respiratory tract. There is significant potential for SARS-CoV-2 and *M.tb* infections to be co-present within individuals and enhance or moderate the respective outcomes of either infection. Here, we investigated how whole blood samples, as well as CD4+ and CD8+ T cells, from individuals hospitalised with acute COVID-19 disease respond to mycobacterial challenge. To do this, samples were assessed by *ex vivo* mycobacterial growth inhibition assays, immune cell phenotyping by mass cytometry, and whole blood cytokine responses to mycobacterial antigens assessed by flow cytometry. These studies identified a subgroup of COVID-19 patients whose blood had an enhanced capacity to inhibit mycobacterial growth. The ability to control mycobacterial growth was associated with the presence of a non *M.tb*-specific CD28+ CD8+ T cell population, with a particular activation status and migratory phenotype. This work improves our understanding of factors involved in mycobacterial control, and may contribute to the design of novel therapies for TB.

## INTRODUCTION

Tuberculosis (TB), caused by *Mycobacterium tuberculosis* (*M.tb*), is the most leading bacterial infectious disease killer worldwide. In 2023 there was a worldwide incidence of 133 per 100,000 persons, with an estimated 10.8 million new cases and 1.25 million deaths from TB^1^. TB is the leading cause of death from a single infectious agent, and the number one killer among HIV-infected individuals^2^. In 2020, the world witnessed the collision of the ancient pandemic caused by *M.tb* with the recent pandemic caused by severe acute respiratory syndrome coronavirus-2 (SARS-CoV-2), the causative agent of coronavirus disease 2019 (COVID-19 disease). The COVID-19 pandemic negatively impacted TB control programmes and reversed the years of progress made in the fight against TB^2^. In the aftermath of COVID-19, fewer people were diagnosed and treated for TB, and for the first time since 2005 the number of cases of TB and associated deaths increased. SARS-CoV-2 and *M.tb* can occupy the same lower respiratory tract niches: LTBI and active TB patients can be infected by SARS-CoV-2 and, similarly, individuals first infected with SARS-CoV-2 can then contract *M.tb*. Both agents have the capacity to alter the local lung environment, orchestrating specific immune responses which could be harmful, neutral or beneficial to a secondary infection^3,4^. There is a very limited understanding of how the immune responses, specifically T cell responses, elicited by *M.tb* and SARS-CoV-2 interact.

This study aimed to provide a better understanding of the host immunological interactions between *M.tb* and SARS-CoV-2 infections. The local immune factors that impact early mycobacterial control, and the possibility to progress to severe disease, are poorly understood and difficult to investigate in clinical settings. Here, whole blood from patients with acute COVID-19 was challenged with mycobacteria, and growth was assessed. Mass and flow cytometry were used for immune cell phenotyping and to investigate *M.tb*-specific T cell cytokine responses.

## RESULTS

### A subset of COVID-19 patients exhibit enhanced control of mycobacterial growth in whole blood assays

The capacity of whole blood from healthy controls and acute COVID-19 patients (Table 1) to inhibit mycobacterial growth was assessed. For this, we performed *ex vivo* Mycobacterial Growth Inhibition Assays (MGIA). Whole blood was incubated with avirulent (BCG) or virulent (H37Rv) mycobacteria (Suppl. Fig. 1A). All healthy controls controlled both mycobacterial species (Fig. 1A and B). Many COVID-19 patients also controlled both infections at a level similar to healthy controls, but there was a subset of individuals who exhibited enhanced mycobacterial control compared to healthy individuals (Fig. 1A and B). Moreover, there was a strong correlation between an individual’s ability to control BCG and H37Rv growth (r=0.62, p=0.0005; Fig. 1C). These data show that a subset of acute COVID-19 patients has enhanced capacity to inhibit mycobacterial growth (subsequently termed ‘controllers, classification based on the median of BCG net growth in whole blood from COVID-19 patients [-0.1506]) (Fig. 1D).

**Fig. 1.**
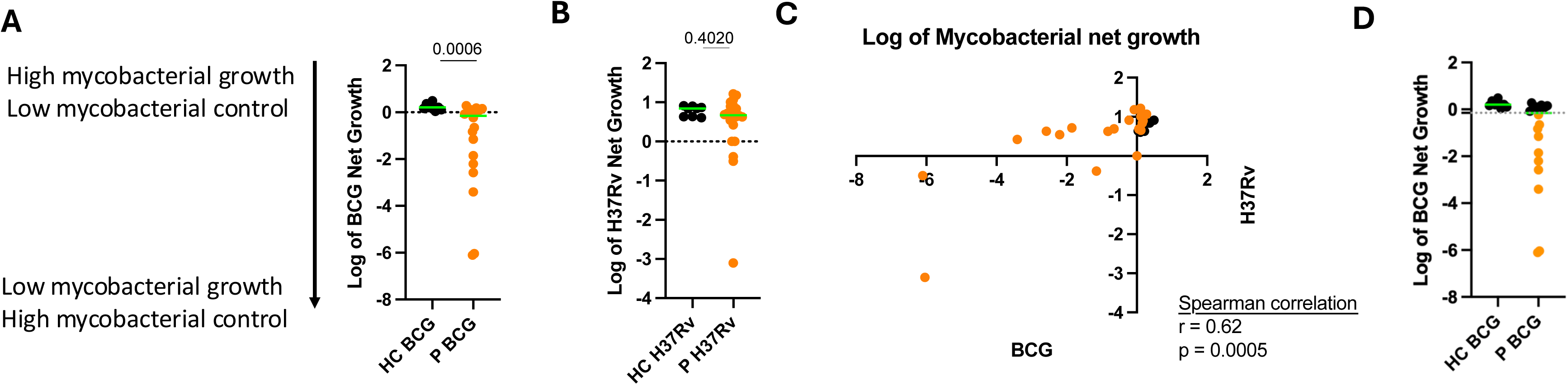
Whole blood of a subset of COVID-19 patients inhibits mycobacterial growth. Log_10_ of BCG **(A)** and H37Rv **(B)** net growth measured using the BACTEC MGIT 960 and whole blood from healthy controls (HC, black, n=9) and acute COVID-19 patients (P, orange, n=20). **(C)** Spearman correlation of Log_10_ of BCG and H37Rv net growth. **(D)** Log_10_ of BCG net growth stratified by median: non-controllers (black, 9 HC and 10 COVID-19 patients) and controllers (orange, n=9 COVID-19 patients).

**Table 1:** Study participants.

| Study No. | Status | Enrolment date | Positive COVID test | Age | Sex | Ethnicity | POB | BCG vaccination | History of TB | Immunosuppression | COVID vaccination | Oxygen |
| --- | --- | --- | --- | --- | --- | --- | --- | --- | --- | --- | --- | --- |
| 100 | Patient | 17-Jan-22 | 16-Jan-22 | 56 | F | White Caucasian | UK | Y | N | N | N | 0 |
| 101 | Patient | 21-Jan-22 | 18-Jan-22 | 72 | F | White Caucasian | UK | Y | N | N | Y (x3) | 0 |
| 102 | Patient | 25-Jan-22 | 22-Jan-22 | 51 | M | White Caucasian | UK | N | N | N | Y (x3) | 3L/min |
| 103 | Patient | 25-Jan-22 | 23-Jan-22 | 64 | F | White Caucasian | UK | Y | N | Y (chemotherapy) | Y (x3) | 0 |
| 104 | Patient | 10-Feb-22 | 09-Feb-22 | 76 | M | White Caucasian | UK | Y | N | N | Y (x3) | 8L/min |
| 105 | Patient | 10-Feb-22 | 09-Feb-22 | 68 | M | White Caucasian | UK | ? | N | N | Y (x3) | 0 |
| 106 | Patient | 24-Feb-22 | 21-Feb-22 | 61 | F | White Caucasian | UK | Y | N | N | Y (x3) | 4L/min |
| 107 | Patient | 24-Feb-22 | 24-Feb-22 | 81 | F | White Caucasian | UK | Y | N | N | Y (x3) | 0 |
| 108 | Patient | 28-Feb-22 | 26-Feb-22 | 59 | M | White Caucasian | USA | N | N | N | Y (x3) | 0 |
| 109 | Patient | 04-Jul-22 | 02-Jul-22 | 53 | M | White Caucasian | UK | ? | N | N | Y (x3) | 0 |
| 110 | Patient | 14-Jul-22 | 13-Jul-22 | 77 | F | White Caucasian | UK | ? | N | N | Y (x3) | 0 |
| 111 | Patient | 18-Jul-22 | 16-Jul-22 | 74 | M | White Caucasian | UK | N | N | N | Y (x3) | 0 |
| 112 | Patient | 21-Jul-22 | 19-Jul-22 | 24 | F | Pakistani | UK | Y | N | N | Y (x2) | 0 |
| 113 | Patient | 14-Jul-22 | 12-Jul-22 | 60 | F | White Caucasian | UK | ? | N | N | Y (x2) | 0 |
| 114 | Patient | 21-Jul-22 | 19-Jul-22 | 88 | M | White Caucasian | UK | Y | N | N | Y (x4) | 0 |
| 115 | Patient | 25-Jul-22 | 24-Jul-22 | 54 | F | White Caucasian | UK | Y | N | N | Y (x3) | 1L/min |
| 116 | Patient | 25-Jul-22 | 23-Jul-22 | 47 | F | White Caucasian | UK | Y | N | N | Y (x3) | 0 |
| 117 | Patient | 28-Jul-22 | 26-Jul-22 | 74 | F | White Caucasian | UK | Y | N | N | Y (x3) | 0 |
| 118 | Patient | 01-Aug-22 | 30-Jul-22 | 86 | F | White Caucasian | UK | Y | N | N | Y (x3) | 2L/min |
| 119 | Patient | 01-Aug-22 | 30-Jul-22 | 89 | M | White Caucasian | UK | ? | N | N | Y (x4) | 0 |
| HC2 | Control | 10-Jan-22 | NA | 25 | F |  |  | N |  | N | Y (x3) | NA |
| HC3 | Control | 21-Jan-22 | NA | 37 | F |  |  | Y |  | N | Y (x3) | NA |
| HC4 | Control | 25-Jan-22 | NA | 47 | M |  |  | Y |  | N | Y (x3) | NA |
| HC5 | Control | 07-Apr-22 | NA | 39 | F |  |  | Y |  | N | Y (x3) | NA |
| HC6 | Control | 07-Apr-22 | NA | 30 | M |  |  | Y |  | N | Y (x3) | NA |
| HC7 | Control | 16-May-22 | NA | 66 | M |  |  | Y |  | N | Y (x3) | NA |
| HC8 | Control | 16-May-22 | NA | 52 | F |  |  | Y |  | N | Y (x3) | NA |
| HC9 | Control | 27-May-22 | NA | 65 | F |  |  | Y |  | N | Y (x3) | NA |

### COVID-19 patients have fewer circulating lymphocytes and more neutrophils and plasmablasts than healthy controls

To investigate if specific immune cell populations were associated with mycobacterial control, we compared the whole blood immunophenotype between COVID-19 mycobacterial controllers and non-controllers using mass cytometry (Fig. 1D). However, because of the small sample size, we first ensured that mass cytometry would allow detection of distinctly abundant cell populations between groups by comparing healthy controls and all COVID-19 patients. We observed that COVID-19 patients had fewer lymphocytes and more neutrophils than healthy controls (Fig. 2A), as previously described^5^. High-dimensionality data of the non-granulocyte population (e.g. all neutrophils excluded; gating strategy in Supplementary Fig. S2) was plotted into a tSNE space, encompassing levels of expression of 36 cellular markers (Supplementary Table S1). As shown in Fig. 2B, the occupied tSNE space was different between the healthy and the COVID-19 groups, indicating distinct cellular compositions. To identify which cellular subsets accounted for these differences, the FlowSOM algorithm was used to distinguish 38 cell populations (as dictated by Phenograph analysis) (Fig. 2C and 2D). Differences in the frequency of these 38 cell populations between the two groups were investigated. Fig. 2E and 2F show an example of a particular cell subset (Population 4) present in the COVID-19 patient group but almost absent in the healthy controls. Population 4 is CD45RO+ CD45RA+ CD19+ CD20-HLA-DR+ CD38+ CD27+ (Fig. 2C), a phenotype consistent with plasmablasts. This same CD38+ CD27+ plasmablast population has been identified as a feature of COVID-19 patients in different patient cohorts, both by flow and mass cytometry^6–8^. These results validated our experimental approach, in which we can identify the distinct cellular composition and marker level expression between the healthy and patient groups by mass cytometry.

**Fig. 2.**
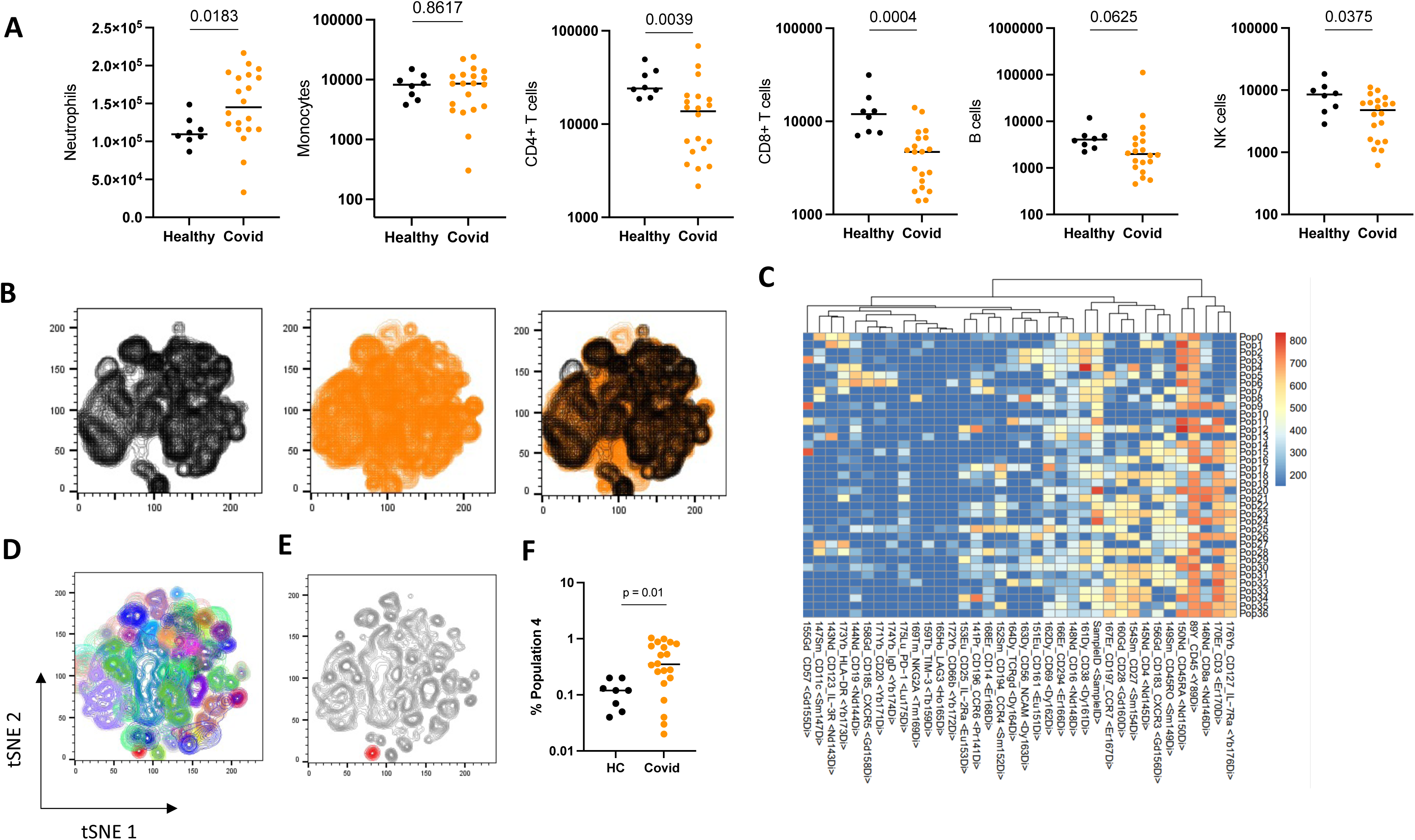
COVID-19 patients present fewer circulating lymphocytes and more neutrophils and plasmablasts than healthy controls. **(A)** Total counts of neutrophils, monocytes, CD4+ T cells, CD8+ T cells, B cells and NK cells present in 270 μl of whole blood from healthy controls (black, n=9) or acute COVID-19 patients (orange, n=20), measured by mass cytometry. **(B)** tSNE space projection of whole blood from healthy controls (black, n=9) and acute COVID-19 patients (orange, n=20), based on the 30-marker Maxpar® Direct™ Immune Profiling Assay™ (MDIPA) and the 6-marker T cell expansion panel 1 (Supplementary Table S1). **(C)** Heatmap depicting levels of expression of the 36 tested markers in each of the 38 immune-cell clusters identified using Phenograph and FlowSOM. **(D)** tSNE space projection of healthy controls and COVID-19 patients showing the 38 cell populations identified using Phenograph and FlowSOM. **(E)** tSNE space projection of healthy controls and COVID-19 patients with Population 4 (corresponding to plasmablasts) highlighted in red. **(F)** Percentage of Population 4 within whole blood of healthy controls (HC, black, n=9) and acute COVID-19 patients (Covid, orange, n=20).

### CD28+ CD8+ T cells are enriched within COVID-19 mycobacterial controllers

The COVID-19 population was divided into two subgroups (Fig. 1D) based on the median of BCG net growth in whole blood from COVID-19 patients (-0.1506): controllers (BCG net growth < median) or non-controllers (BCG net growth > median) (Fig. 3A). The high-dimensional data within a tSNE space was plotted, revealing differences between groups (controllers vs non-controllers). The same 38 distinct cell populations identified in Fig. 2C and 2D were applied to the new tSNE space (Fig. 3B, 3C and 3D). Two T cell populations with distinct frequencies in the controller and non-controller groups were identified (Fig. 3E and 3F). A TCRγ8+ CD57+ T cell population was present at a higher frequency in the non-controller group (Fig. 3F) but did not correlate with mycobacterial control (Fig. 3G).

**Fig. 3.**
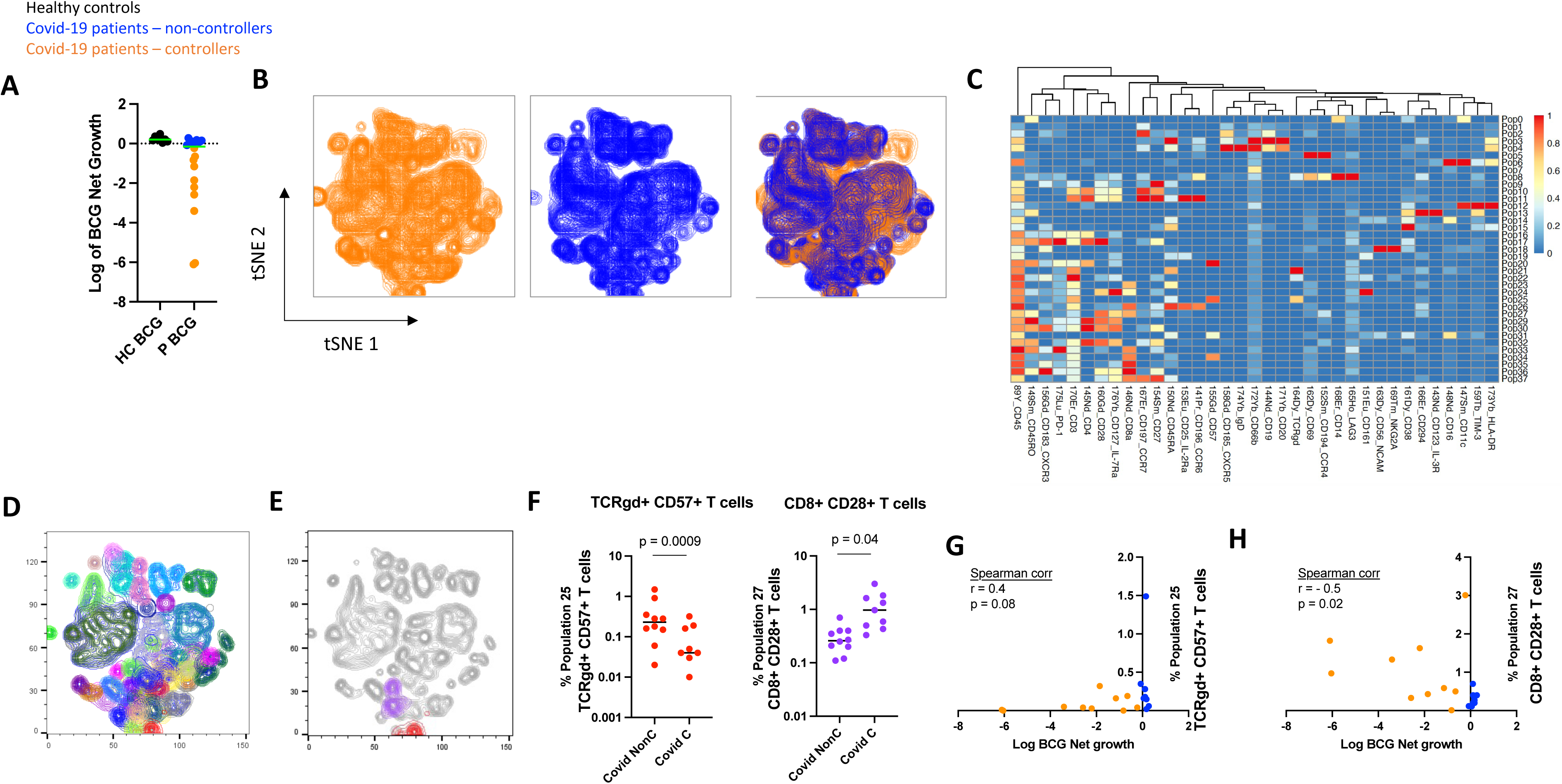
COVID-19 controllers present high levels of a CD28+ CD8+ T cell subset which correlates with mycobacterial growth inhibition. (A) The COVID-19 patient (P) population was divided in two groups depending on their ability to control BCG mycobacterial growth (cut-off: median), measured using whole blood in the BACTEC MGIT 960 (see Fig. 1). Non-controller group (blue, n=10); controller group (orange, n=10). (B) tSNE space projection of whole blood from acute COVID-19 patients, controllers (orange, n=10) and non-controllers (blue, n=10), based on the 30-marker Maxpar® Direct™ Immune Profiling Assay™ (MDIPA) and the 6-marker T cell expansion panel 1. (C) Heatmap depicting levels of expression of the 36 tested markers in each of the 38 immune-cell clusters identified using Phenograph and FlowSOM. (D) tSNE space projection of acute COVID-19 patients showing the 38 cell populations identified using Phenograph and FlowSOM. (E) tSNE space projection of acute COVID-19 patients with highlighted Population 25 (red) and Population 27 (purple). (F) Percentages of Populations 25 (TCRγδ+ CD57+ T cells) and 27 (CD8+ CD28+ T cells) within whole blood of non-controller (NonC) and controller (C) acute COVID-19 patients. Spearman correlation between Log_10_ BCG net growth and the whole blood percentages of Populations 25 (G) and 27 (H).

However, a specific memory CD127+ (IL7Ra) CD8+ CD28+ T cell subset was more abundant in the controller group (Fig. 3F) and negatively correlated with mycobacterial growth (Fig. 3G), suggesting a potential role in mycobacterial control.

### ESAT-6/CFP-10- and PPD-specific T cell antigen responses do not correlate with mycobacterial control

Having identified a CD8+ T cell subset associated with mycobacterial control (Fig. 3G), we investigated whether the capacity to inhibit mycobacterial growth was due to the presence of mycobacteria-specific CD8+ and CD4+ T cells. *In vitro* stimulation of whole blood from COVID-19 patients with mycobacterial ESAT-6/CFP-10 proteins and Purified protein derivative (PPD) was performed (see gating strategy in Supplementary Figure 3). The capacity of both CD8+ and CD4+ T cells from COVID-19 patients to secrete cytokines in response to non-specific cell activators (i.e. PMA) was assessed as a control (Fig. 4A, 4B and 4C). Increased production of IFNγ and TNF by CD8+ T cells from COVID-19 patients after ESAT-6/CFP-10 protein stimulation was observed, but not in the healthy controls (Fig. 4A, 4B and 4C). The same pattern was detected in CD4+ T cells for IFNγ, IL-2 and TNF, and in CD8+ and CD4+ T cells for TNF upon PPD stimulation (Fig. 4A, 4B and 4C). There was a strong correlation between IFNγ production by CD4+ and CD8+ T cells after ESAT-6/CFP-10 protein stimulation (Fig. 4D, r=0.82, p<0.0001) but not between CD8+ T cell IFNγ production after ESAT-6/CFP-10 full protein and PMA/Ionomycin stimulation (Fig. 4E, r=-0.007, p=0.98), confirming that the same individuals are responding to full ESAT-6 and CFP-10 proteins in CD4+ and CD8+ T cells. The magnitude of IFNγ responses in CD8+ T cells after ESAT-6/CFP-10 full protein or PPD did not correlate with inhibition of mycobacterial growth (Fig. 4F and 4G), suggesting that the mechanism underlying mycobacterial control is not T-cell ESAT-6/CFP-10-specific. Thus, although an increase in cytokine secretion (e.g. TNF) following ESAT-6/CFP-10 or PPD stimulation was observed in COVID-19 patients compared to healthy controls, this was not responsible for the mycobacterial control detected in the blood of a subset of patients.

**Fig. 4.**
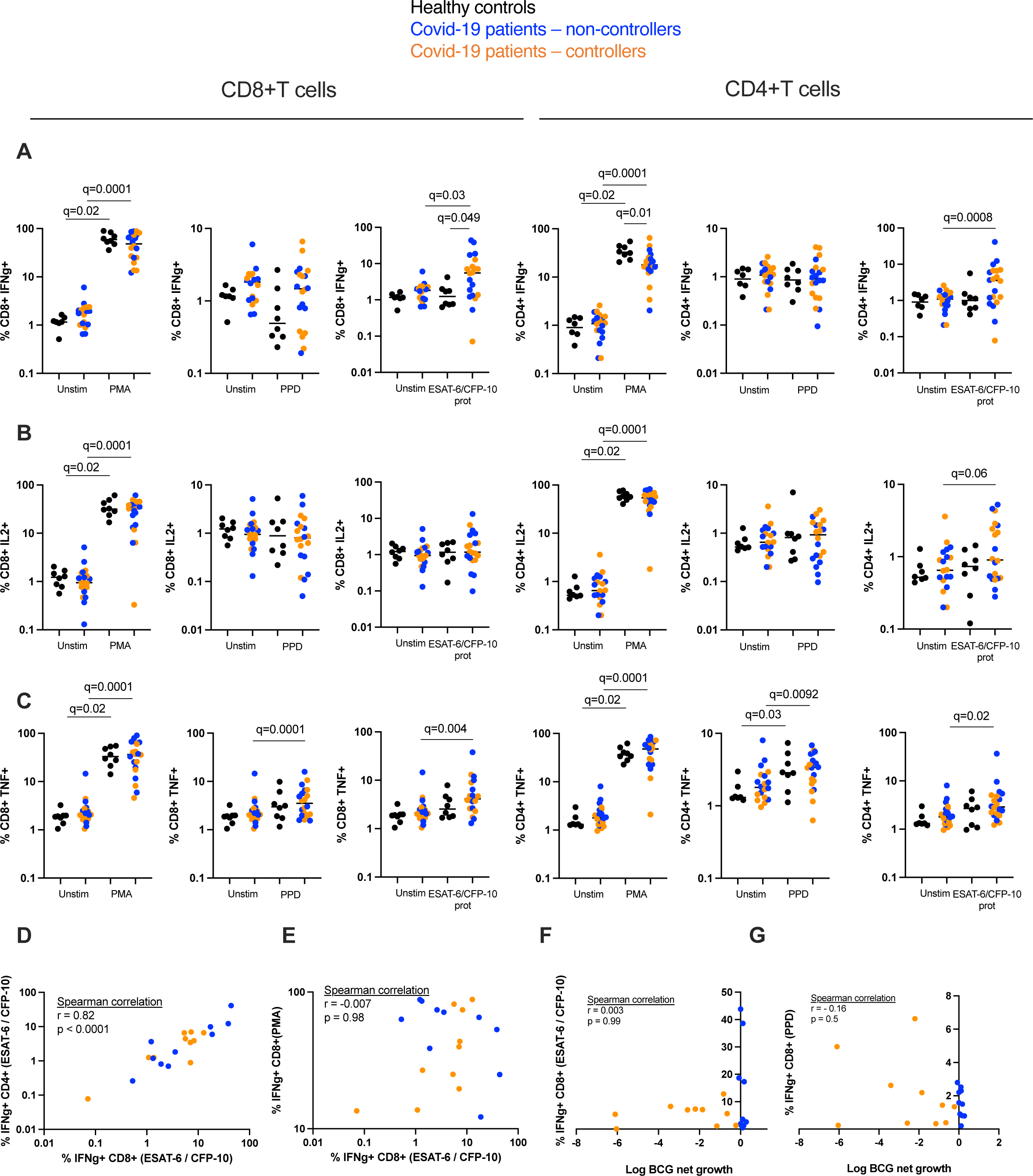
A subset of COVID-19 patients’ CD4+ T cells recognise *Mycobacterium tuberculosis* antigens, but this does not correlate with mycobacterial control. Percentage of whole blood CD8+ and CD4+ IFNγ+ **(A)**, IL2+ **(B)** and TNF+ **(C)** T cells measured by flow cytometry after stimulation with PMA/Ionomycin, PPD or ESAT-6/CFP-10 full proteins, from healthy individuals (black, n=9) and acute COVID-19 patients (controllers in orange n=10, non-controllers in blue n=10). Spearman correlation between % IFNγ+ CD8+ and % **(D)** IFNγ+ CD4+ T cells after stimulation with ESAT-6/CFP-10 full proteins, and **(E)** IFNγ+ CD8+ T cells after PMA/Ionomycin stimulation, from acute COVID-19 patients (n=20). Spearman correlation between % IFNγ+ CD8+ **(F)** T cells after stimulation with ESAT-6/CFP-10 full proteins and % log_10_ BCG net growth, from acute COVID-19 patients’ whole blood (n=20).

### The CD28+ CD8+ T cell population associated with mycobacterial control possesses a unique activation and migratory capacity phenotype

To further characterise the CD28+ CD8+ T cell population associated with mycobacterial control among severe acute COVD-19 patients, levels of T cell activation, senescence and chemokine receptors were assessed. Expression of specific activation markers and chemokine receptors within the CD28+ CD8+ T cell population did not differ between the controller and the non-controller groups (Fig. 5A). Correlations between specific activation markers and chemokine receptors (the ones with the highest/detectable levels of expression) were analysed, (Fig. 5B) both for the CD28+ and CD28-CD8+ T cell populations. The CD28+ population showed strong correlations between specific activation markers and chemokine receptors. For instance, HLA-DR and PD-1 expression negatively correlated with CCR7, a receptor responsible for directing cell migration towards secondary lymphoid organs. Conversely, CD57 and CD69 expression levels positively correlated with CXCR3, receptor responsible for guiding T cells to the site of inflammation. These correlations were absent in the CD28-CD8+ T cell population. These results suggest that the CD28+ CD8+ T cell population associated with mycobacterial control has a specific activation status and migratory capacity, potentially resulting in higher numbers of activated cells at the site of infection.

**Fig. 5.**
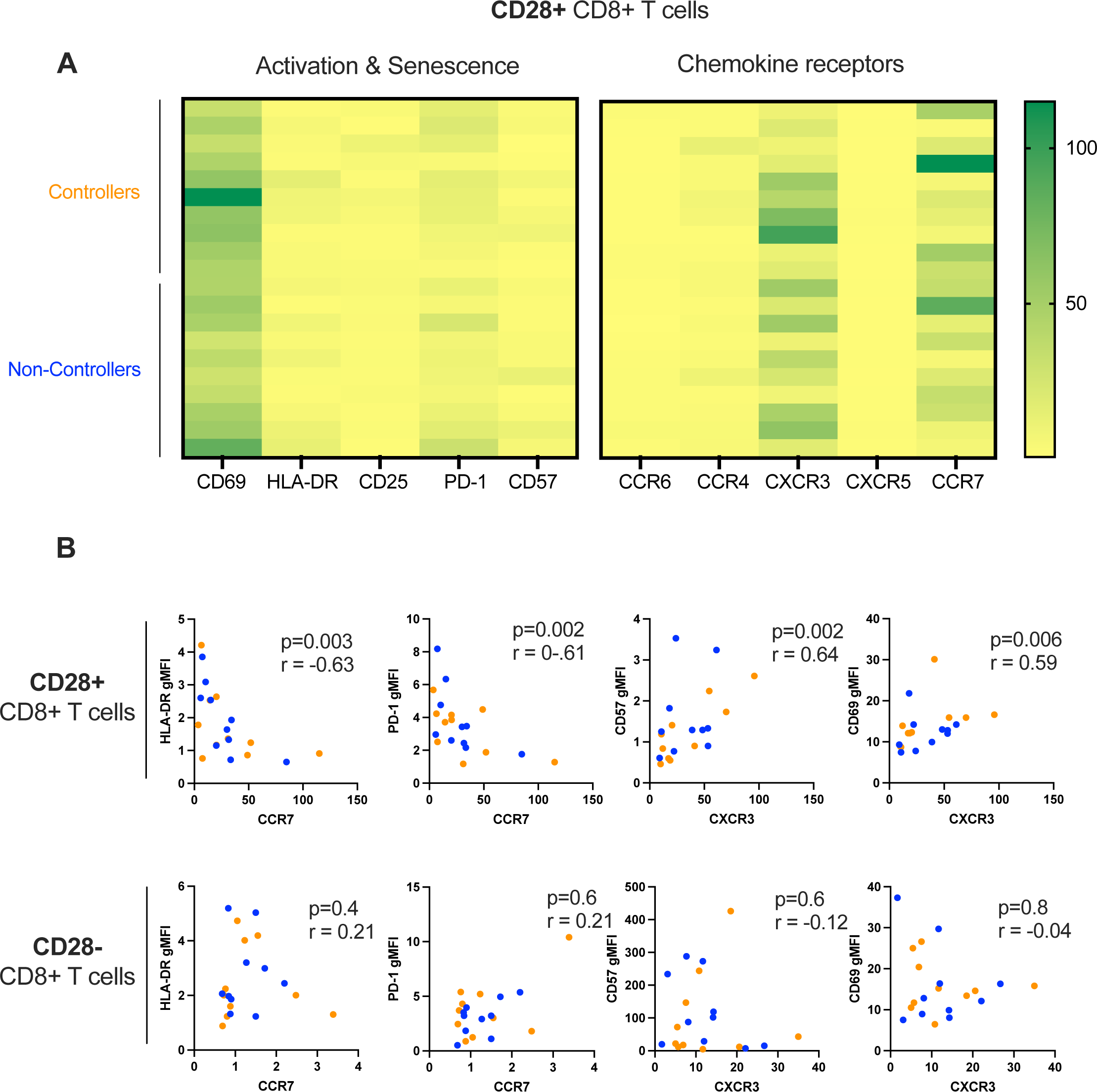
Characterisation of the CD28+ CD8+ T cell population associated with mycobacterial control. **(A)** Heatmap depicting levels of expression of activation and senescence markers, as well as of chemokine receptors, of the CD28+ CD8+ T cell population associated with mycobacterial control, measured by mass cytometry. Expression is shown for both mycobacterial controllers and non-controllers of the COVID-19 group. **(B)** Spearman correlations of expression of activation markers (HLA-DR, PD-1, CD57 and CD69) and chemokine receptors (CCR7 and CXCR3) in both CD28+ and CD28-CD8+ T cell populations with the COVID-19 group (controllers in orange n=10, non-controllers in blue n=10).

## DISCUSSION

Here, we describe the immunological interactions between mycobacteria and samples from patients admitted to hospital with severe acute SARS-CoV-2 infection. The capacity of H37Rv or BCG to grow within whole blood was measured using a mycobacterial growth inhibition assay^9^. This assay provides a comprehensive, unbiased functional readout which includes a variety of possible immune mechanisms and their complex interactions. The whole blood of a subset of acute COVID-19 patients showed improved ability to restrict mycobacterial growth (i.e. controllers). The heterogeneity of responses was greater among COVID-19 patients compared to controls which was particularly evident with BCG (which is biologically plausible given the differential growth between BCG and H37Rv^9^). The strong correlation between BCG and H37Rv net growth provides useful validation of the functional output from this assay.

Using mass cytometry in whole blood, an increased subset of memory CD28+ CD8+ T cells in COVID-19 patients was identified, (Fig. 3F) which negatively correlated with BCG net growth (Fig. 3H).

CD8+ cytokine responses to ESAT-6/CFP-10 proteins and PPD did not correlate with mycobacterial control (Fig. 4F). This suggests that the identified memory CD8+ T cell population targets different epitopes than those contained within the ESAT-6/CFP-10 proteins and PPD, or that the mechanism behind this effect is not mycobacteria-specific. Antigen-specific CD8+ T cells exert protective functions in TB ^10^, and specific T cell receptor repertoires have been linked to disease control and progression after *M.tb* infection^11^. Further investigations are required to explore the antigen-specificity of this particular T cell subset. It is also possible that it belongs to an unconventional T cell population similar to Mucosal-Associated Invariant T (MAIT) cells ^12^. This unconventional, MR1-restricted T cell population has been associated with anti-mycobacterial responses^13–15^ . Furthermore, a TRAV1-2+ CD8+ T cell subset with MAIT or MAIT-like features has been found to be enriched in the airways of active TB patients^16^. Recently, CD8+ T cells were shown to be crucial for early control of *M.tb* in macaques^17^. Specifically, circulating CD8+ CD28+ T cells are diminished in patients with active TB compared to LTBI^18,19^. In contrast, pulmonary TB patients (PTB) have more CD8+ CD28-T cells than healthy controls, and this specific T cell subset is positively associated with disease progression and sputum bacillary loads^20^. Here, we show CD8+ CD28+ T cells are associated with an ex vivo mycobacterial control phenotype. Furthemore, increased CD8+CD57+ T cells has been reported in TB patients compared with age-matched healthy tuberculin skin test-positive donors^21^, and with healthy controls^22^. CD57+ *M.tb*-specific CD4+ and CD8+ T cells are more abundant in active TB than in LTBI^23^. ^24^ and to healthy controls^25^. We have shown an increased abundance of a TCRγδ+ CD57+ T cells in the non-controller group, compared to mycobacterial controllers.

Decreased CD28 and increased CD57 expression on T cells has been described in aging^26,27^, and during persistent antigen exposure and may impair T cell effector functions (reviewed in Strioga *et al.*^28^). Using a functional assay in the context of acute SARS-CoV-2 infection we have identified increased (TCRγδ+ CD57+) and decreased (CD8+ CD28+) T cells in individuals unable to control mycobacterial growth. We speculate that this model stimulated an inflammatory environment which enabled such differentiation and we show the power of combining whole blood MGIT assays with mass cytometry and downstream computational analysis to identify new functional links between specific immune cell populations and mycobacterial control.

It has been reported that both active TB and previously treated TB enhance the severity and mortality of COVID-19^34–37^. Interestingly, mouse models of infection have suggested a protective role for underlying *M.tb* infection to acute SARS-CoV-2 disease^38,39^. *In vitro*, supernatants from PBMCs infected with *M.tb* decreased levels of SARS-CoV-2 replication in Calu-3 human airway epithelial cells^39^. In an observational, cross-sectional study, plasma concentrations of cytokines associated with systemic inflammation were lower in TB patients previously infected with SARS-CoV-2 infection, than in TB patients without prior acute COVID-19^40^. Both CD4+ and CD8+ T cells primed *in vitro* with BCG-derived peptides showed increased reactivity to specific SARS-CoV2 derived peptides^41^. *In silico* studies using machine-learning algorithms have suggested TCR cross-reactivity between SARS-CoV-2 and *M.tb* (shared peptide antigens between both pathogens, presented through specific TCRs)^42^.

Contrary to the concept that pre-existing acute lung infection might universally increase host susceptibility to further pulmonary pathogens, our findings suggest heterogeneity and that acute COVID-19 may confer, in some cases, some enhanced control of mycobacterial infections. This might help explain why increased rates of *M.tb* infection have not been reported within COVID-19 patients. In a recent study in mouse, the lung inflammatory microenvironment at the time of infection facilitated innate immune control of SARS-CoV-2 replication^33^, providing experimental evidence that distinct, pre-existing pulmonary conditions may impact innate immune response and the outcome of new infections. A deeper understanding of the underlying mechanisms might provide new insights into novel therapies for mycobacterial diseases.

In summary, we report that the whole blood of a subset of acute COVID-19 patients has the ability to decrease mycobacterial growth. We have identified a specific CD28+ CD8+ memory T cell population associated with enhanced mycobacterial control. These findings might help better understand the dynamics of the TB and SARS-CoV-2 pandemics, as well as provide background knowledge for the identification of new therapeutic opportunities for mycobacterial disease.

## MATERIALS AND METHODS

### Ethics, consent and samples

The study was approved by the Health Research Authority and Health and Care Research Wales (IRAS project ID 303500, Protocol number RG_21-128, REC Reference 21/EM/0211) and all aspects of the study were conducted according to the principles of the Declaration of Helsinki and Good Clinical Practice. Adult patients (≥18 years) receiving care at University Hospitals Birmingham NHS Foundation Trust (Birmingham, UK), with a new diagnosis of COVID-19 (defined as a positive SARS-COV-2 PCR result within the preceding 72 hours), were invited to participate. Healthy controls were University of Birmingham staff recruited under local ethical approval (REC Reference ERN_17-0411). All volunteers provided written informed consent prior to sample collection. Whole blood was collected in BD Vacutainer Lithium Heparin tubes (and used within 1 hour of collection in ex vivo assays). Participant characteristics are shown in Table 1.

### MGIT Mycobacterial growth inhibition assay

A whole blood mycobacterial growth inhibition assay was used as previously described^43^. Briefly, duplicate tubes containing 300 μl of whole blood were mixed with 300 μl of RPMI-MGIT media containing RPMI-1640 Medium HEPES modification with 25mM HEPES (Sigma-Aldrich), 2mM L-glutamine (Sigma-Aldrich) and 10% pooled human AB serum (Sigma-Aldrich). The blood-media mix was then seeded with ∼150 colony forming units (cfu) BCG Pasteur or *M.tb* H37Rv and treated with either 10 μM Dexamethasone (Sigma-Aldrich), 10 ng/ml Tocilizumab (ThermoFischer Scientific) or untreated, and incubated on a 360° rotator at 37°C for 96 hours (volume of the mycobacterial stock was calculated to give a time to positivity, TTP, of 6.5 days, previously determined to give optimal differential responses). Sterile water was then used to lyse the cells, and the lysate was transferred to a BACTEC MGIT tube supplemented with PANTA antibiotics and OADC enrichment broth (Becton Dickinson, UK). Tubes were transferred to the BACTEC 960 (Becton Dickinson, UK) and incubated at 37°C until detection of positivity by fluorescence (TTP, hours). On day 0, duplicate direct-to-MGIT viability control tubes were prepared by inoculating supplemented MGIT tubes with ∼150 cfu of mycobacteria. The TTP was converted to Log_10_ cfu using stock standard curves of TTP against inoculum volume and cfu. The net growth ratio was calculated as Log_10_(sample cfu/control cfu). A smaller net growth value indicates less bacillary replication and therefore represents greater mycobacterial control.

### Whole blood stimulation

500 μl of whole blood was aliquoted into 2ml sterile tubes and left unstimulated or stimulated with one of: PMA/ionomycin (eBioscience™ Cell Stimulation Cocktail); 10 μg/ml of ESAT-6 and CFP-10 full proteins (BEI Resources) + anti-human CD28/CD49d antibodies at 1 μg/ml (BD Bioscience). Samples were incubated at 37°C and 5 μg/ml of Brefeldin A (Sigma-Aldrich) was added after 5 hours. Samples were incubated at 37°C for a further 14-16 hr.

EDTA (2mM, ThermoFischer Scientific) was added to all samples and incubated at room temperature for 15 min. Red blood cells were lysed using FACS lysing solution (BioLegend). Samples were centrifuged at 800g for 7 min and the supernatant discarded. The pellet was washed with PBS, centrifuged at 800g for 7 min and the supernatant discarded. 1ml of freezing media (FCS [Biosera] with 10% DMSO [ThermoFischer Scientific]) was added to each sample and transfer into a cryovial for storage at -80°C until use.

### T cell Cytokine production by flow cytometry

Cryopreserved samples were thawed, immediately diluted in 10ml of RPMI (Sigma-Aldrich), centrifuged at 800g for 7 min and the supernatant discarded. Pellets were resuspended in 150 μl of PBS, transferred to 96-well V-bottom plates and centrifuged (800g, 3 min, 4°C).

Wells were washed with PBS and centrifuged again. Fc-receptor blocking was performed following manufacturer’s instructions (Miltenyi Biotec). Cells were washed twice with PBS and centrifuged (800 g, 3min) before performing surface staining for 20 min at 4°C in the dark with anti-human CD3-CF594 antibody (ThermoFischer Scientific), anti-human CD4-BV605 antibody (Becton Dickinson) and anti-human CD8-BV510 antibody (BioLegend). Cells were washed twice with PBS and centrifuged before fixing/permeabilising following manufacturer’s instructions (eBioscience). Intracellular staining was performed using anti-human IFNγ-AF700 antibody (Becton Dickinson), anti-human TNF-PE-Cy7 antibody (ThermoFischer Scientific) and anti-human IL-2-FITC antibody (ThermoFischer Scientific).

Cells were washed twice and acquired using the BD LSR Fortessa II. Analysis was performed using FlowJo v10.8.1 and GraphPad Prism 9. An example of the gating strategy is outlined in Supplementary Fig. S8.

### Immune cell phenotyping by mass cytometry

Whole blood was collected in sodium heparinised tubes (BD Vacutainer). Heparin (Sigma-Aldrich) was added to 300 μl of blood for a final concentration of 100 U/ml and incubated for 20 min at room temperature. 270 μl of heparin-blocked blood was transferred into one 5 ml tube containing the dry antibody pellet for the Maxpar Direct Immune Profiling Assay (MDIPA, Standard Biotools). The antibodies of the Maxpar® Direct™ T Cell Expansion Panel 1 (Standard Biotools, Supplementary Table S1) were added, and tubes incubated for 30 min at RT. 250 μl of Cal-Lyse lysing solution (ThermoFischer Scientific) was added to each tube and incubated in the dark for 10 min at room temperature. 3ml of Maxpar Water (Standard Biotools) was added to each tube and incubated for 10min at RT. Tubes were centrifuged at 300g for 5 min and washed three times with Cell Staining Buffer (CSB). Cells were fixed in 1ml of 1.6% formaldehyde solution (Standard Biotools) for 10 min at RT and centrifuged at 800g for 5min, discarding the supernatant. Cells were stained with 1ml of Cell-ID Intercalator-Ir (Standard Biotools) into Maxpar Fix and Perm Buffer (Standard Biotools) to a final concentration of 125 nM for 1h at RT. Cells were centrifuged at 800g for 5 min, resuspended in 1ml of freezing media (FCS [Biosera] with 10% DMSO [ThermoFischer Scientific]) and transferred into a cryovial for storage at -80°C until analysis with the Helios (CyTOF) mass cytometer.

Preliminary analysis was performed using the Maxpar Pathsetter software. Fcs files were analysed with FlowJo version 10.8.1 and R version 4.1.2. Live singlet cells were selected and granulocytes excluded using the gating strategy presented in Supplementary Fig. S8. 10,000 events from the “live singlets no granulocytes” population were down sampled from each file using the DownSample (version 3.3.1) plugin, and concatenated into one single fcs file for high dimensionality reduction analysis. To estimate the number of clusters (or distinct cell populations) present in the combined dataset, the plugin PhenoGraph (version 3.0) was run using default parameters, and identified 38 cell populations. The plugin FlowSOM was then used to identify the 38 cell populations within the dataset using default parameters, and setting the number of meta clusters as 38. Cell population frequencies for each sample were extracted using FlowJo, and compared across cohorts using a non-parametric Mann-Whitney test. Correlation between cell population frequencies and the MGIT assays were performed using the non-parametric Spearman’s correlation test. The geometric mean fluorescence intensity (gMFI) of each marker within each of the 38 cell populations (across all samples) was extracted using FlowJo and plotted using R, after min-max standardized normalization.

### Statistical Analysis

Non-parametric T-tests were performed when comparing two groups, paired if the two groups corresponded to the same individuals (Wilcoxon), unpaired if the groups corresponded to different individuals (Mann-Whitney), and corrected for multiple comparisons.

## ACKNOWLEDGEMENTS

We are grateful to all study participants, the wider research team who supported this study from University Hospitals Birmingham NHS Foundation Trust and the University of Birmingham, and to the Flow Cytometry platform at the University of Birmingham.

## AUTHOR CONTRIBUTIONS

MKO and AC conceived the study, obtained funding and provided overall guidance. AL, HS, JP and MKO designed and performed experiments, analysed and interpreted data. JGB analysed mass cytometry data. CJ, HH, SF and ZS performed additional experiments. MD and CAG were clinical study co-ordinators. EW, HK and ES undertook patient recruitment and sample collection. AL, MKO and AC prepared the manuscript. All authors contributed to manuscript revision, read and approved the submitted version.

## DECLARATION OF INTERESTS

We declare no competing interests.

## FUNDING

Funding was provided by UKRI MRC (Grant Ref: MR/W015374/1).

**Supplementary Fig. S1. Mycobacterial growth inhibition assay.** Workflow of the MGIT mycobacterial inhibition assay.

**Supplementary Fig. S2. Example of mass cytometry gating strategy.** Cells were gated on single cells, intact singlets, alive cells, CD45+ cells and non-granulocytes (CD16-CD66b-) for downstream analysis.

**Supplementary Fig. S3. Example of flow cytometry gating strategy.** Cells were gated on lymphocytes, single cells, alive cells, CD3+ cells and CD4+ or CD8+ cells **(A)**, before looking at their levels of IFNγ, IL2 or TNF cytokine production **(B)**.

## Notes

### Competing Interest Statement

The authors have declared no competing interest.

