## Supplementary figures and images for "Differential control of mycobacteria among COVID-19 patients is associated with CD28+ CD8+ T cells"

### Supplementary Figures

**Supplementary Figure 1**

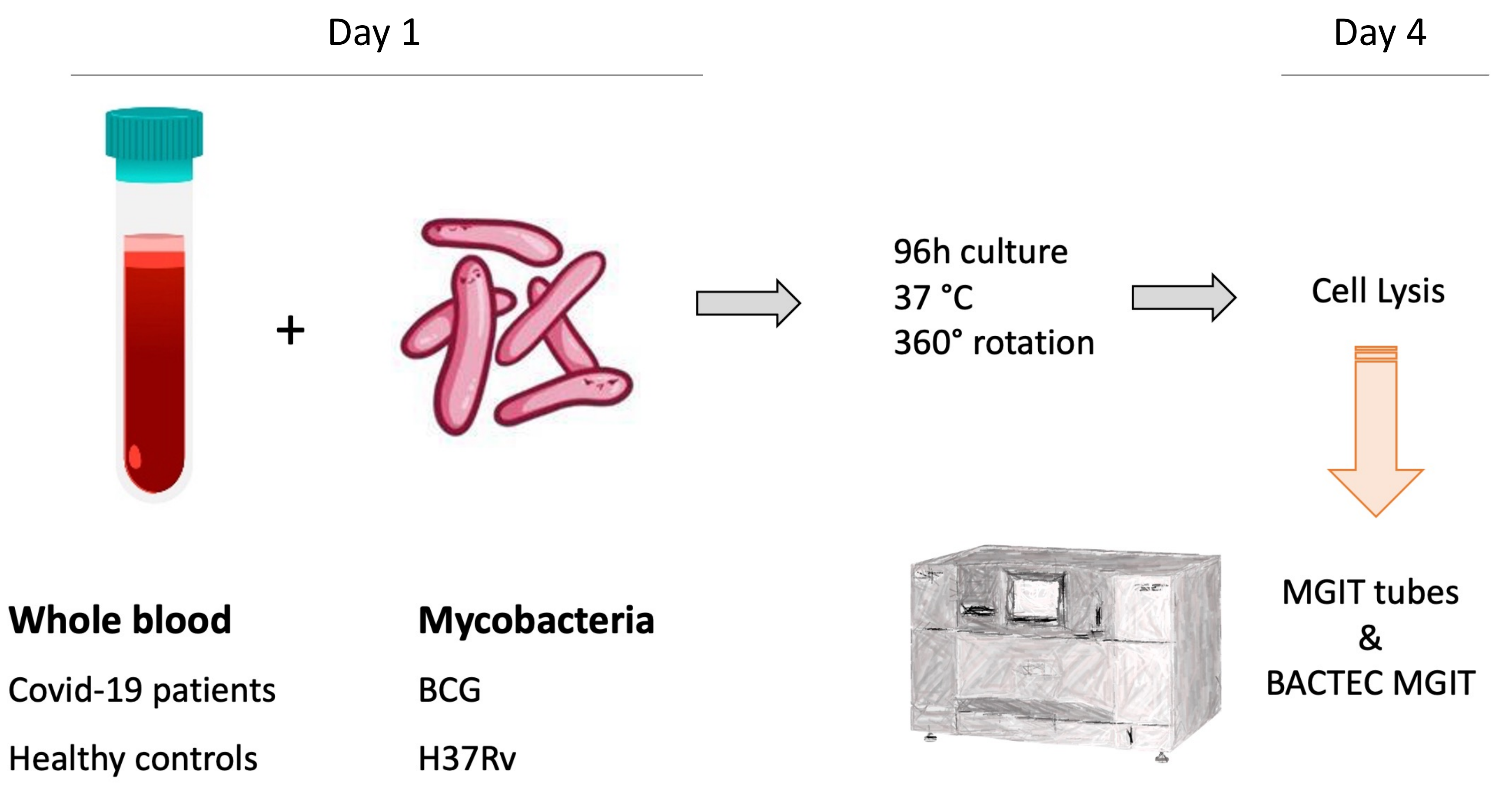

#### Supplementary Figure 2

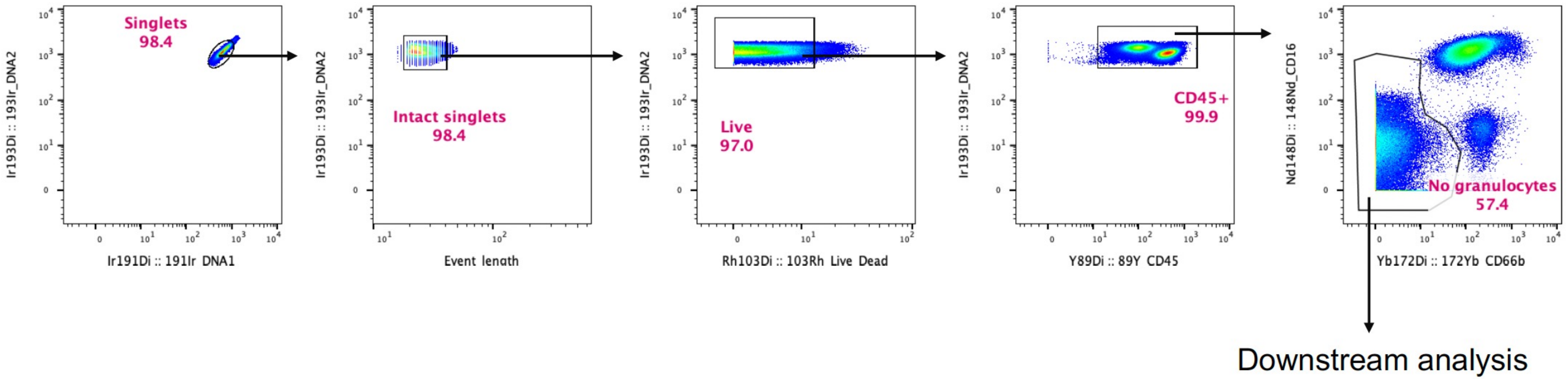

#### Supplementary Figure 3

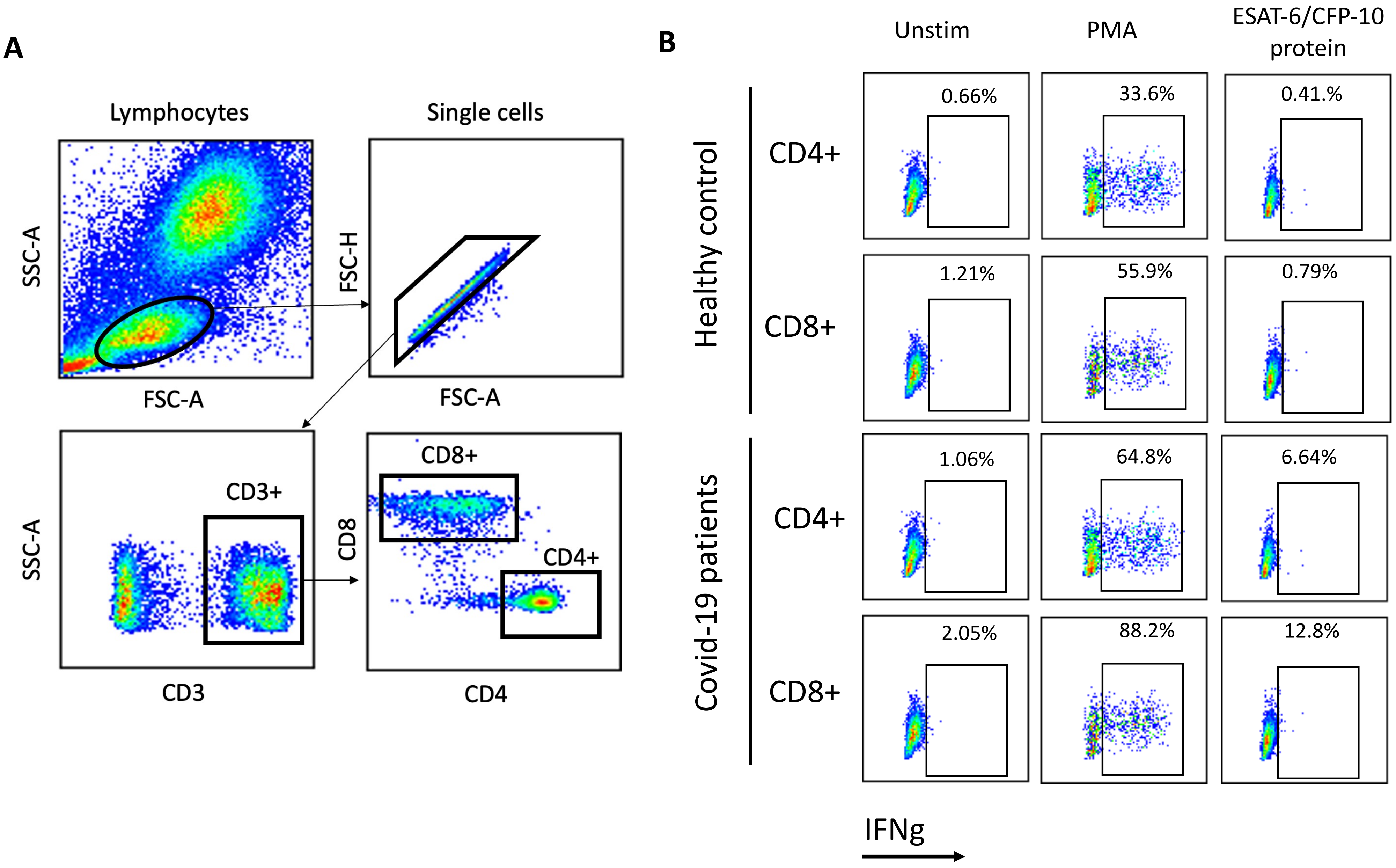
