## Supplementary table 1 for "Differential control of mycobacteria among COVID-19 patients is associated with CD28+ CD8+ T cells"

**Supplementary Table S1.**

| Panel | Metal - Label | Target |
|---|---|---|
| MDIPA | 142Ce | Bead |
| | 140Ce | Bead |
| | 191Ir | DNA1 |
| | 153Eu | CD25_IL-2Ra |
| | 175Lu | Bead |
| | 165Ho | Bead |
| | 151Eu | CD161 |
| | 193Ir | DNA2 |
| | 103Rh | Live_Dead |
| | 156Gd | CD183_CXCR3 |
| | 150Nd | CD45RA |
| | 145Nd | CD4 |
| | 144Nd | CD19 |
| | 154Sm | CD27 |
| | 152Sm | CD194_CCR4 |
| | 160Gd | CD28 |
| | 158Gd | CD185_CXCR5 |
| | 155Gd | CD57 |
| | 164Dy | TCRgd |
| | 170Er | CD3 |
| | 166Er | CD294 |
| | 168Er | CD14 |
| | 176Yb | CD127_IL-7Ra |
| | 172Yb | CD66b |
| | 174Yb | IgD |
| | 171Yb | CD20 |
| | 147Sm | CD11c |
| | 146Nd | CD8a |
| | 161Dy | CD38 |
| | 167Er | CD197_CCR7 |
| | 141Pr | CD196_CCR6 |
| | 89Y | CD45 |
| | 148Nd | CD16 |
| | 163Dy | CD56_NCAM |
| | 173Yb | HLA-DR |
| | 143Nd | CD123_IL-3R |
| | 149Sm | CD45RO |
| T cell expansion panel 1 | 159Tb | CD366_Tim3 |
| | 162Dy | CD69 |
| | 165Ho | CD223_LAG3 |
| | 169Tm | CD159a_NKG2A |
| | 175Lu | CD279_PD-1 |
| | 209Bi | TIGIT |
